# Planarians employ diverse and dynamic stem cell microenvironments to support whole-body regeneration

**DOI:** 10.1101/2022.03.20.485025

**Authors:** Blair W. Benham-Pyle, Frederick G. Mann, Carolyn E. Brewster, Enya R. Dewars, Dung M. Vuu, Stephanie H. Nowotarski, Carlos Guerrero-Hernández, Seth Malloy, Kate E. Hall, Lucinda E. Maddera, Shiyuan Chen, Jason A. Morrison, Sean A. McKinney, Brian D. Slaughter, Anoja Perera, Alejandro Sánchez Alvarado

**Author notes:** Correspondence should be addressed to Alejandro Sánchez Alvarado. These authors contributed equally and author order was determined by coin flip (Blair W. Benham-Pyle, Frederick G. Mann).

## Abstract

Stem cells enable regeneration by self-renewing and differentiating as instructed by a local microenvironment called a niche^1–3^. In most cases, the repair or replacement of tissues is fueled by tissue-specific or lineage-restricted stem cells that proliferate in response to local injury and apoptosis^4–11^. However, in organisms that regenerate using abundant adult pluripotent stem cells, the stem cell niches that support tissue repair have not been identified or characterized. Since these adult pluripotent stem cells are often more widely distributed and plentiful than lineage-restricted stem cells of other organisms, defining their microenvironments may uncover alternative forms of stem cell regulation^12–14^. Here we used unbiased spatial transcriptomics to define the cellular and molecular environments that support pluripotency in the highly regenerative freshwater planarian *Schmidtea mediterranea*. We determined that stem cells associate with a diverse collection of differentiated cell types, and these associations are highly dynamic during regeneration. We explored associations with two distinct cell types: secretory cells we term ‘hecatonoblasts,’ and intestinal cells. While both cell types regulate stem cell proliferation, their spatial relationships to stem cells defy the concept of a single regenerative niche. Thus, the planarian stem cell pool is likely maintained by a dynamic collection of distinct microenvironments that cooperatively power whole-body regeneration.

## MAIN TEXT

The freshwater planarian *Schmidtea mediterranea* possesses an abundant population of adult pluripotent stem cells that allow it to regenerate its entire body from a tiny fragment^12–14^. Yet, little is known about the local environments that support stem cells during regeneration. We sought to identify cell types and molecules associated with pluripotent stem cells during regeneration by taking advantage of emerging tools in spatial transcriptomics. Slide-seqV2, an unbiased spatial transcriptomic method, captures mRNA transcripts from tissue sections onto barcoded beads with known X/Y positions^15, 16^. Importantly, Slide-seqV2 beads are 10 microns in diameter, allowing us to capture transcripts of a highly localized cellular environment containing 1-5 cells. We reasoned that when combined with existing single-cell RNA-seq data of regenerating planarians, a spatial atlas with this resolution could allow us to identify the cell types closely associated with stem cells and contributing to regenerative competence (Figure 1a)^17^.

**Fig. 1:**
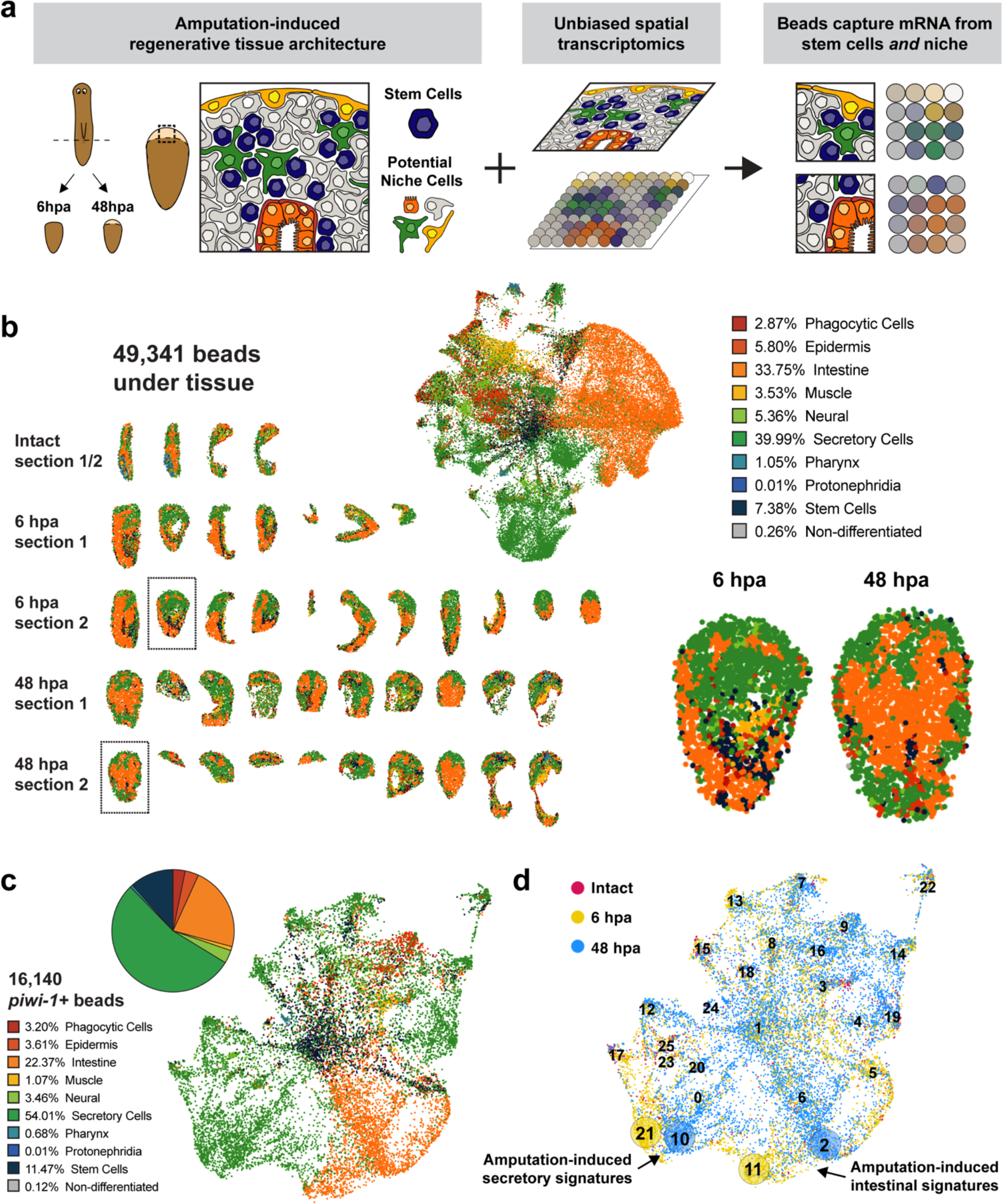
Slide-seqV2 identifies cell types that dominate stem cell microenvironments during regeneration. **a,** Diagram of experiment. Left, regenerating planarian fragments were collected at 6 and 48 hours post regeneration (hpa) to analyze cellular organization during regeneration. Middle, approximately 10 fragments were co-embedded around an intact animal, and sectioned for Slide-seqV2. Two adjacent sections were analyzed on separate pucks. Right, Slide-seqV2 beads co-capture mRNA from stem cells and their neighbors. **b,** Tissue signatures captured in the data. Left, ‘Row embedding’ of beads under the regenerating tissue fragments, colored by strongest tissue annotation produced by Seurat’s Label Transfer function from a previous scRNA-seq dataset (Benham-Pyle et al., 2021). Boxed fragments are enlarged at right. Middle, UMAP plot of complete dataset colored by tissue annotation. Percentages indicate the number of beads of each tissue annotation. Right, example fragments from 6 and 48 hpa. **c,** UMAP plot of *piwi-1^+^* bead subset colored by tissue annotation. Frequency of tissue annotations is presented both as percentages and summarized in a pie chart. **d,** UMAP plot of *piwi-1+* bead subset colored by timepoint. Cluster numbers are superimposed on the plot; clusters of interest are indicated with a circle around the numbers.

To generate Slide-seqV2 data, planarians were amputated, and regenerating tissue assayed at 6 and 48 hours post amputation (hpa) (Figure S1a). Live regenerating fragments were arranged around an intact animal with wounds facing inward in OCT embedding media, frozen, and sectioned for either Slide-seqV2, histological staining, or nuclear staining (Figure S1b). Slide-seqV2 captured significantly more mRNA transcripts under planarian tissue, which facilitated the removal of background beads from the dataset, the definition of individual tissue sections, and the reorientation of fragments along their anterior-posterior axis (Figure S1c-h). We analyzed data from 49,341 under-tissue beads and used Seurat’s LabelTransfer function to predict which tissue types had contributed to each bead (Figure 1b, Figure S1i-k)^18^. *In vivo* expression patterns of tissue-specific transcripts were recapitulated by Slide-seqV2, albeit with lower resolution, thereby validating our dataset (Figure S2, Supplementary Table 1).

We next sought to identify the cells and tissues likely to contribute to regenerative stem cell niches. We hypothesized that cell types in proximity to stem cells would co-deposit their mRNA onto the same beads. Therefore, we focused our analysis on only the beads that captured *piwi-1*, a well-established marker for planarian stem cells^13^. Of the 49,341 beads under tissue, 16,140 (32.7%) captured at least one *piwi-1* transcript (Figure S3a). Re-clustering and analysis of *piwi-1*^+^ beads revealed unexpected heterogeneity in stem cell microenvironments (Figure S3b). Only 11% of *piwi-1^+^* beads were dominated by a stem cell signature, while most beads extensively captured mRNA from other cell types (Figure 1c, Figure S3c-f). The majority of beads were dominated by gene signatures enriched in secretory cells (54.01%). Planarian secretory cells are a family sometimes referred to as ‘parenchymal cells’ that have diverse expression patterns across the animal but have not previously been linked to stem cell function^17, 19, 20^. The second most frequent tissue signature was intestinal (22.37%). Prior studies have found that planarian stem cells are intercalated between gut branches, and that intestinal genes can regulate stem cell proliferation and differentiation^21–23^.

Given the heterogeneity of stem cell microenvironments predicted by the dataset, we sought to verify their spatial and temporal distribution. For each of the *piwi-1^+^* bead clusters, we calculated the proportion of beads from 6 or 48 hpa, as well as the average distance of the beads from the wound (Figure 1d, Figure S4a-d). While some clusters were evenly composed of beads from 6 or 48 hpa, several were dominated by beads from one timepoint (Figure S4a-b). Moreover, most clusters appeared at different distances from the wound at 6 or 48 hpa (Figure S4d). Visualization of markers of select bead clusters validated the spatial and temporal biases identified by the spatial atlas (Figure S4e-m). Together, our results indicate that planarian stem cell microenvironments are diverse and highly dynamic during regeneration. To characterize and functionally test some of the identified stem cell microenvironments, we focused on bead clusters with intestinal and secretory signatures, given their abundance in the *piwi- 1^+^* subset.

The spatial atlas predicts dynamic associations between stem cells and intestinal cells during regeneration, with distinct molecular and cellular components at 6 and 48 hpa (Figure 2a-b). To test this model, we visualized intestinal cells and stem cells *in vivo* in intact animals and in regenerating animals at 6 and 48 hpa. As expected based on prior literature, stem cells can be found near intestinal cells at all timepoints, often intercalated between gut branches (Figure 2c, Figure S5a,c,e)^21^. We also imaged intestinal and stem cells, along with the mitotic marker phospho-Histone H3 (H3P) using high-resolution confocal microscopy to better understand the spatial relationship between intestinal cells and proliferative stem cells at 6 and 48 hpa (Figure 2d-e). We observed stem cells in the vicinity of the intestine, but rarely were the two cell types in direct contact, regardless of condition. To quantify the positions of these cells in a rigorous manner, we used the artificial intelligence program CellPose to classify and localize 337,935 cells in the images (Figure S6, Figure S7)^24, 25^. We found that *piwi- 1^+^*/H3P^+^ proliferating stem cells were enriched 10-40 microns away from the intestinal cells, but depleted in the area immediately adjacent to them (Figure 2f, Supplementary Table 2). This enrichment was only detected in intact animals and at 6 hpa, but not at 48 hpa. To understand if this enrichment was specific to dividing stem cells, we also quantitated overall stem cell enrichment in these regions. We found similar enrichment of *piwi-1^+^* stem cells 10-40 microns away from the intestine, at all three timepoints (Figure 2g). Notably, stem cell enrichment was highest at 6 hpa. In general, we did not find a consistent relationship between proximity to intestinal cells and rate of stem cell proliferation (Figure 2h). These experiments reveal a stem cell ‘goldilocks zone’ 10-40 microns away from the intestine where stem cells accumulate and proliferate, particularly at 6 hpa. The discovery of a consistent gap between intestinal cells and stem cells likely indicates signaling between the two cell types is contact independent.

**Fig. 2:**
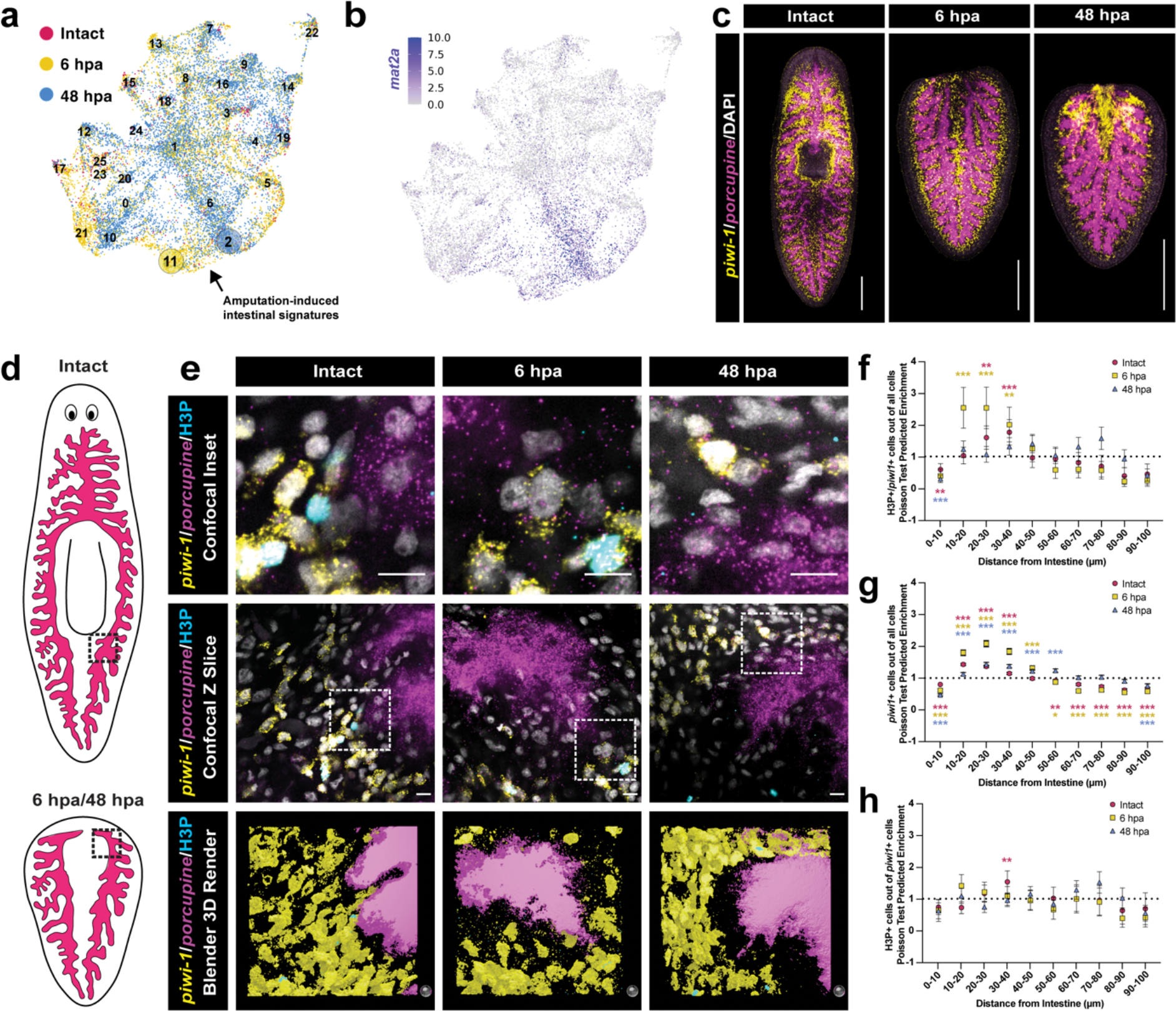
Stem cells are dynamically enriched at different distances from intestinal cells. **a,** UMAP plot of *piwi-1^+^* bead subset colored by timepoint. Intestinal clusters of interest are indicated. **b,** UMAP plot of *piwi-1*^+^ bead subset colored by the number of UMI for the intestinal marker *mat2b*. **c,** Double fluorescent *in situ* hybridization of the intestinal marker *porcupine* (magenta), *piwi-1* (yellow), and DAPI (white) in a single confocal Z- section. Scale bars = 500 microns. **d,** Cartoon of planarian intestine. Region of interest indicated with a dotted box. **e,** First row, insets showing detail of a single confocal slice of double fluorescent *in situ* hybridization of *porcupine* (magenta) and *piwi-1* (yellow), immunofluorescence of H3P (cyan), and DAPI (white). Scale bars = 10 microns. Second row, wider area image of First row for context. Scale bars = 10 microns. Third row, 3- dimensional (3D) rendering of 40 micron deep confocal image stacks encompassing single slices shown in rows above. Scale sphere in lower right= 10 microns in diameter. **f,** Plot showing fraction of all cells that are H3P^+^/*piwi-1*^+^ as a function of distance from the intestine. **g,** Plot showing fraction of all cells that are *piwi-1*^+^ as a function of distance from the intestine. **h,** Plot showing fraction of *piwi-1*^+^ cells that are H3P^+^/*piwi-1*^+^ as a function of distance from the intestine. For f,g,h, cells and distances were identified and quantified by the program CellPose. Analysis was performed for images from intact animals (red circles), 6 hpa (yellow squares), and 48 hpa (blue triangles). Data points represent means +/- SE. Asterisks indicate statistical significance * = p < .01, ** p < 0.001, *** p < 0.0001. All p-values presented in Supplementary Table 2.

We next explored the spatial relationship between secretory cells and stem cells.

Among the many *piwi-1^+^* bead clusters dominated by secretory cell signatures, we chose to focus on the microenvironments represented by clusters 17 and 21. Both of these clusters were enriched at 6 hpa and mutually marked by the secretory gene *matrix metalloproteinase-1* (*mmp-1*), among other genes with similar functions like *mmp-2* and *tolloid-like-1* (Figure 3a-b). These two clusters were also among the nearest to the wound. To test these predictions, we visualized *mmp-1* and *piwi-1* in regenerating fragments and found that both transcripts were present near the wound, but expressed in different cell types (Figure 3c, Figure S5b,d,f). We found *mmp-1^+^* secretory cells sparsely arranged in a ringlike pattern in the mesenchyme around the pharynx of the animal. They were roughly 3 times as wide as *piwi-1^+^* stem cells and irregularly shaped with prominent projections. While *mmp-1^+^* secretory cells have previously been observed, they have not been extensively characterized beyond capture in single-cell RNA-seq datasets (Fincher *et al.*, ‘Parenchymal’ Subcluster 9; Benham-Pyle *et al.*, ‘Parenchymal’ Subcluster 18)^17, 19^. While the term ‘parapharyngeal’ has been used to denote cells in this region of the animal, we have chosen to name the *mmp-1^+^* cell population to distinguish it from similarly distributed cell populations, such as *foxA^+^* stem cells and pharyngeal precursors, as well as other secretory cell populations^17, 19^.

**Fig. 3:**
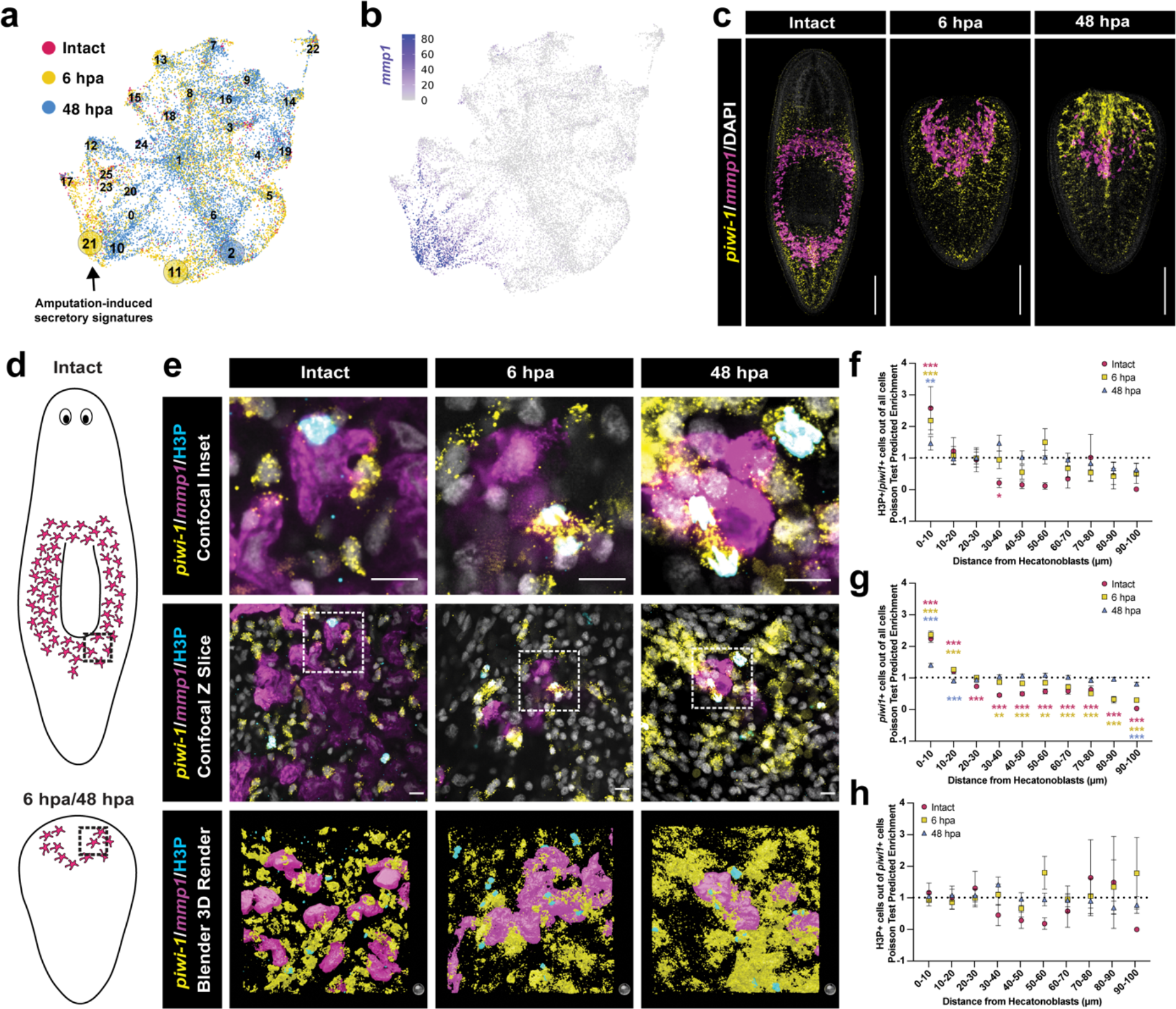
Stem cells stably and directly contact hecatonoblasts. **a,** UMAP plot of *piwi-1^+^* bead subset colored by timepoint. Secretory cell clusters of interest are indicated. **b,** UMAP plot of *piwi-1*^+^ bead subset colored by the number of UMI for the secretory cell marker *mmp-1*. **c,** Double fluorescent *in situ* hybridization of the secretory marker *mmp-1* (magenta), *piwi-1* (yellow), and DAPI (white) in a single confocal Z-section. Scale bars = 500 microns. **d,** Cartoon of hecatonoblasts in planarians. Region of interest indicated with a dotted box. **e,** First row, insets showing detail of a single confocal slice of double fluorescent *in situ* hybridization of *mmp-1* (magenta) and *piwi-1* (yellow), immunofluorescence of H3P (cyan), and DAPI (white). Scale bars = 10 microns. Second row, wider area image of First row for context. Scale bars = 10 microns. Third row, 3-dimensional (3D) rendering of 40 micron deep confocal image stacks encompassing single slices shown in rows above. Scale sphere in lower right= 10 microns in diameter. **f,** Plot showing fraction of all cells that are H3P^+^/*piwi-1*^+^ as a function of distance from the intestine. **g,** Plot showing fraction of all cells that are *piwi-1*^+^ as a function of distance from the intestine. **h,** Plot showing fraction of *piwi-1*^+^ cells that are H3P^+^/*piwi-1*^+^ as a function of distance from the intestine. For f,g,h, cells and distances were identified and quantified by the program CellPose. Analysis was performed for images from intact animals (red circles), 6 hpa (yellow squares), and 48 hpa (blue triangles). Data points represent means +/- SE. Asterisks indicate statistical significance * = p < .01, ** p < 0.001, *** p < 0.0001. All p-values presented in Supplementary Table 2.

Because these large cells have processes making direct contact with the stem cells (Figure 3e), we name these cells *‘hecatonoblasts’* after hecatoncheires, the many- handed giants from Greek mythology.

As we did for the intestine, we next sought to define the spatial relationship between stem cells and hecatonoblasts during regeneration. We again collected high- resolution confocal microscopy images of *mmp-1^+^* hecatonoblasts and *piwi-1^+^* stem cells along with the mitotic marker H3P. We observed close associations of stem cells and hecatonoblasts, with the two cell types often in direct contact (Figure 3d,e). In some cases hecatonoblasts appeared to be wrapped around stem cells. This was observed at all three timepoints. We classified and localized the positions of 280,372 cells using CellPose to analyze the rate of proliferation as a function of proximity to hecatonoblasts. In contrast to what we observed for the intestine, the abundance of *piwi-1^+^*/H3P^+^ proliferating stem cells was significantly higher in the area immediately adjacent to hecatonoblasts (Figure 3f. We observed a similar trend for all *piwi-1^+^* cells, meaning that we again did not observe a difference in proliferation rate as a function of proximity to hecatonoblasts (Figure 3g-h). These experiments identified a close association between stem cells and hecatonoblasts, creating the potential for direct cell-cell communication.

After finding spatial associations of stem cells with intestinal cells and hecatonoblasts, we sought to determine if these cell types regulate stem cell function during regeneration. We identified genes enriched in the bead clusters that co-captured *piwi-1* and either intestinal or hecatonoblast markers, then selected 23 from each for functional testing (Supplementary Tables 3-4). We first used *in situ* hybridization to visualize the expression patterns of all genes in both sets, then compared these patterns to our existing single-cell RNA-seq atlas^17^. Genes co-captured with *piwi-1* and intestinal transcripts were strongly enriched in intestinal cells in the scRNA-seq data (Figure S8a). Accordingly, nearly all probes yielded characteristic intestinal expression patterns *in vivo* (Figure S8b). For genes co-captured with *piwi-1* and hecatonoblast transcripts, we generally found secretory cell-like expression patterns, with some transcripts also being detected in other tissues, such as the stem cells, central nervous system, and muscle (Figure S9a). The expression patterns we observed *in vivo* were consistent with expression data from scRNA-seq (Figure S9b).

Having determined the tissue specificity of all the gene targets, we next set out to test their roles in regulating stem cell proliferation via RNA interference. For our initial screen, we tested the roles of these genes in intact animals and regeneration. We visualized the stem cell marker *piwi-1* and the mitotic cell markers phospho-Histone H3 (H3P) in intact animals 7 days post feeding, and during the wound-induced burst of proliferation at 48 hpa (Figure 4a). Significantly, genes enriched in both cell types had effects on wound-induced proliferation, but little effect on intact animals (Figure S10a). We identified five intestinal and nine secretory genes that altered proliferating stem cell densities by at least 25% with a p-value of less than p = .01 (Figure 4b-c). Of these, two intestinal genes and four secretory genes resulted in regeneration or survival defects (Figure 4d, FigureS10b-d). *Innexin* (SMED30010974) and *tubulin alpha* (SMED30021235) are specifically expressed in the intestine and yielded similar RNAi phenotypes, including marked reductions in the number of mitotic nuclei at 48 hpa and occasional failure to survive or regenerate after amputation. *Tetraspanin-18a* (SMED30012544) and *mmp-1* (SMED30019930) are expressed predominantly in hecatonoblasts, and their knockdown yielded mild decreases in numbers of mitoses and occasional failure to survive amputation. The genes *tubulin beta* (SMED30016265) and *cytoplasmic dynein* (SMED30008568) are expressed in hecatonoblasts, but also in neurons, muscle, and stem cells. Their knockdown resulted in severe stem cell, proliferation, survival, and regeneration defects. Together, these phenotypes indicate that hecatonoblasts and intestinal cells not only associate with stem cells but also express genes that regulate their function. Thus, unbiased spatial transcriptomics was able to identify multiple distinct stem cell microenvironments that contribute to whole body regeneration.

**Fig. 4:**
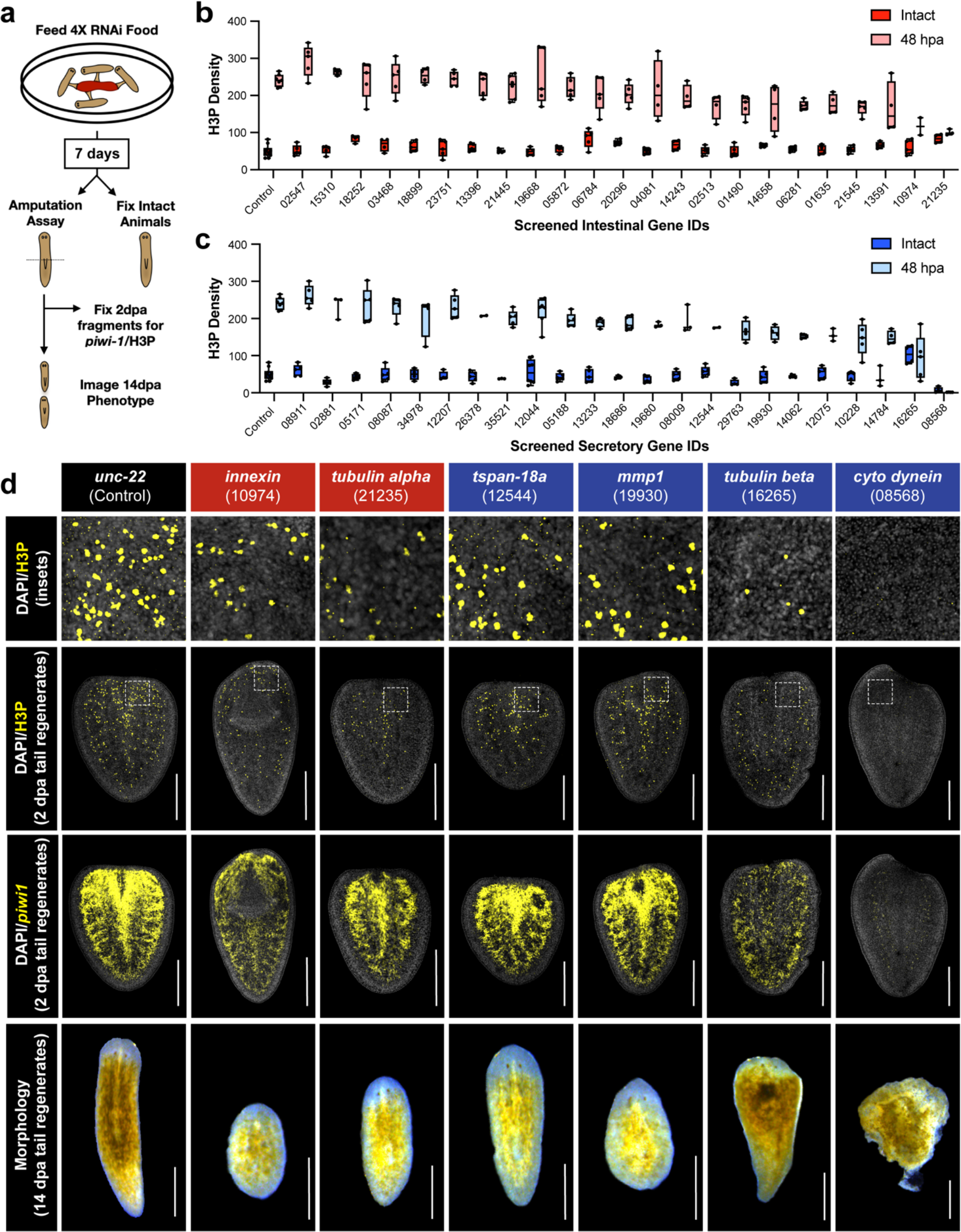
Genes from hecatonoblast and intestinal beads regulate stem cell proliferation. **a,** Schematic of RNAi experiment. Animals were fed RNAi food, and either fixed or amputated 7 days after the final feeding. **b,** Plot depicts number of H3P^+^ nuclei/mm^2^ in intact and 48 hpa animals following RNAi of 24 genes found on *piwi^+^* intestinal beads. Center line, median; box limits, upper and lower quartiles; whiskers, minimum and maximum. Leading “SMED300” was dropped from gene names to save space. **c,** Plot depicts number of H3P^+^ nuclei/mm^2^ in intact and 48 hpa animals following RNAi of 24 genes found on *piwi^+^* secretory beads. **d,** RNAi phenotypes of intestinal/hecatonoblast genes. Top row: detail of confocal max projections showing H3P (yellow) and DAPI (white) at 48 hpa for notable RNAi conditions. Gene names boxed in red are intestinal; those boxed in blue are secretory. Second row: Confocal max projections showing H3P and DAPI throughout regenerating planarian fragments. Boxed areas indicate regions shown in top row. Third row: Confocal max projections showing *piwi-1* (yellow) and DAPI (white) throughout regenerating planarian fragments. Bottom row: Morphological phenotypes at 14 dpa from regenerated tail fragments. Scale bars = 500 microns.

## DISCUSSION

Planarian stem cells display numerous properties which are inconsistent with a spatially restricted niche. In most well studied niches, adult stem cell function is regulated by a collection of nearby differentiated cells^26–35^. In each of the cited examples, the niche maintains a population of resident stem cells devoted to producing the cell types that comprise that tissue. Cells that make up the niche are typically in direct contact with adult stem cells and directly regulate their potency and proliferation. As a result, differentiation is generally initiated when stem cells lose contact with the niche. In contrast, it is unlikely that planarian stem cells are committed to a single niche or tissue lineage. Planarian stem cells can migrate across great distances to regenerate missing tissues^36^. Single-cell sequencing of planarian stem cells reveals a large, distributed, and heterogeneous population of *piwi-1^+^* cells with distinct subsets expressing lineage specific markers^37^. In addition, coarsely-defined spatial regions are enriched for particular tissue progenitors, such as *ovo^+^* or *foxA^+^* stem cell progenitors^38, 39^. However, the expression of lineage markers does not preclude a stem cell from producing daughter cells of a different lineage, and expression of lineage- specific markers may be stochastic or linked to the cell cycle^40^. Most importantly, it appears that spatially-defined lineage bias is reset during the process of whole-body regeneration^41^. Given these observations, the planarian adult stem cell pool would require either a large and distributed niche or an alternative strategy for regulating fate and potency.

To investigate these possibilities, we exploited spatial transcriptomics and published single cell sequencing data to identify the cellular components of planarian stem cell niches. We found no unifying major tissue component of stem cell microenvironments. Instead, we found that stem cells interact with a diverse collection of differentiated cells and these interactions are responsive to injury (Figure 1).

Disruption of just one of the many stem cell microenvironments reduces regenerative capacity (Figure 4). Therefore, our results indicate that planarian regenerative capacity may depend on diversity and plasticity in both the stem cells and their microenvironments. These results may help explain why even stem cell-rich planarian fragments are incompetent to regenerate when below a certain size^17^. While these fragments have hundreds of stem cells, they may not have the necessary biomass to recreate the diversity of microenvironments required for regeneration competence. If indeed the microenvironments produced during regeneration are just as important as the stem cells themselves, future work characterizing each of them and their roles in stem cell regulation will prove critical to understanding planarians’ exceptional regenerative capacity. Emerging evidence indicates that mammalian adult stem cell niches can transiently alter their function in response to injury, and that mammalian stem cells retain potency after exiting their canonical niches^1, 4, 42^. Therefore, it is possible that mammalian stem cells, like planarian stem cells, rely on diverse and distributed microenvironments to respond to injury and that exploration of these microenvironments might reveal conserved principles of tissue repair.

## MATERIALS AND METHODS

### Animal husbandry

Asexual *S. mediterranea* planarians (strain CIW-4 or “C4”) were cultured in recirculating culture system as described previously^43^. Tissue biopsies were taken from animals that had been starved for at least one week. Tail fragments were isolated by using 1.25 mm biopsy punches. Fragment size was standardized by positioning the punch over the tip of the animals’ tails. The isolated tissue comprised a fragment of roughly the most posterior 1.25 mm tissue of the animal. Small animals used for *in situ* hybridization were starved for at least two weeks prior to use.

### Tissue handling and sectioning

Fresh planarian tissue was received in 1x PBS and fragments were embedded together in OCT by timepoint. Tissue fragments were sunk in OCT and arranged in a circle (10- 15 fragments oriented closely together with an overall diameter of ∼3 mm) around an intact animal. Tissue blocks were then flash frozen at −70°C (HistoChill, Novec™ 7000). After freezing, tissue blocks acclimated to −13°C in a cryostat (Thermo, CryoStar NX70) for 30 minutes prior to sectioning. Tissue blocks were then mounted on a cutting block with OCT and sectioned at a 5° cutting angle with 10 μm section thickness. Pucks were held to clean glass via surface tension (DiH20) and tissue sections mounted directly on each puck. Pucks were then removed from the cryostat and placed into a 1.5 mL Eppendorf tube containing hybridization buffer and prepared for library preparation.

Pre- and post-puck sections were also collected on glass slides for each tissue block sectioned. These sections were air-dried at RT for 20 minutes, fixed in 4% PFA for 15 minutes, and then slides were stained; half the slides with hematoxylin/eosin and half with fluorescent DAPI (1:500). Slides were coverslipped and submitted for imaging. The remaining tissue was wrapped in aluminum foil and returned to −70°C and stored for processing at a later date.

### Puck synthesis, library preparation and sequencing

The Slide-seqV2 pucks used in this study were generated and sequenced at the Broad Institute (Cambridge, MA) by Dr. Fei Chen’s group according to the methods and supplementary information provided in the *Nature Biotechnology* publication “Highly sensitive spatial transcriptomics at near-cellular resolution with Slide-seqV2”^16^.The pucks were received at the Stowers Institute for Medical Research on small glass coverslips in 1.7 mL LoBind tubes at room temperature and were stored in the dark at 4°C. The spatial barcode sequencing file of the pucks were generated via monobase ligation chemistry and provided by Dr. Chen’s group to be used for later analysis.

Following mounting, pucks with tissue adhered were immediately immersed in 200 μL of hybridization buffer (6X SSC, 2 unit/μL Lucigen RNase inhibitor) for 27 minutes at room temperature to facilitate the binding of mRNA to the spatially barcoded beads of the puck. First strand synthesis was performed in RT solution (115 μL water, 40 μL 5X Maxima RT buffer, 20 μL 10 mM dNTPs, 5 μL RNase Inhibitor, 10 μL 50 μM Template Switch Oligo, 10 μL Maxima H-RTase) for 30 minutes at room temperature followed by hours at 52°C. After reverse transcription, the tissue was removed by adding 200 μL 2X Tissue Digestion Buffer (100 mM Tris pH 8.0, 200 mM NaCl, 2% SDS, 5 mM EDTA, 32 unit/μL Proteinase K) and incubation at 37°C for 30 minutes.Following this incubation, 200 μL of Wash Buffer (10 mM Tris pH 8.0, 1 mM EDTA, 0.01% Tween-20) was then added and the mixture pipetted to remove the beads from the coverslip. A series of three washes were performed on the beads using 200 μL Wash Buffer for the first two washes and 200 μL 10 mM Tris-HCl (pH 8.0) for the last wash and by pelleting for 3 minutes at 3000 RCF, removing the supernatant and resuspending. Samples were then treated with exonuclease I solution (170 μL water, 20 μL ExoI buffer, 10 μL ExoI) for 50 minutes at 37°C. A series of two washes were performed on the beads using 200 μL Wash Buffer and pelleting for 3 minutes at 3000 RCF, removing the supernatant and resuspending. Samples were then resuspended in 0.1 N NaOH and incubated for 5 minutes at room temperature followed by a series of two washes in 200 μL Wash

Buffer.Beads were resuspended in 200 μL TE, pelleted, supernatant removed and finally resuspended in Second Strand Mix (133 μL water, 40 μL 5X Maxima RT buffer, 20 μL 10 mM dNTPs, 2 μL 1 mM dN-SMRT oligo, 5 μL Klenow Enzyme) and incubated for 1 hour at 37°C. The beads were washed three times with 200 μL Wash Buffer then resuspended with 200 μL water, pelleted for 3 minutes at 3000 RCF and finally resuspended in cDNA PCR mix (88 μL water, 100 μL Terra PCR Direct Buffer, 4 μL Terra Polymerase, 4 μL 100 μM Truseq PCR handle primer, 4 μL 100 μM SMART PCR primer). cDNA was amplified by PCR using the following program:

● 98°C, 3 minutes
● 4 cycles of:

○ 98°C, 20 seconds
○ 65°C, 45 seconds
○ 72°C, 3 minutes
● 9 cycles of:

○ 98°C, 20 seconds
○ 67°C, 20 seconds
○ 72°C, 3 minutes
● 72°C, 5 minutes
● 4°C, forever

The PCR product was purified with 0.6X AMPure XP beads (Beckman Coulter, A63881) twice, resuspended in 20 μL water, and checked for quality and quantity using a Bioanalyzer (Agilent) and Qubit Fluorometer (ThermoFisher).

Subsequent library preparation was performed according to manufacturer’s directions for the Nextera XT kit (Illumina, FC-131-1096) starting with 600 pg of cDNA and using a specific P5-Truseq PCR hybrid oligo in place of the Nextera XT i5 adapter (15ul Nextera PCR mix, 8 μL water, 1 μL 10 μM P5-Truseq PCR hybrid oligo, 1 μL 10μM Nextera N70X oligo).The resulting short fragment libraries were checked for quality and quantity using the Bioanalyzer (Agilent) and Qubit Fluorometer (ThermoFisher).Sequencing was performed on two High-Output flow cells of an Illumina NextSeq 500 instrument using NextSeq Control Software 2.2.0.4 with the following paired read lengths: 42 bp read 1, 8 bp I7 index, and 42 bp read 2.

### Data processing and alignment

Following sequencing, Illumina Primary Analysis version NextSeq RTA 2.4.11 and Secondary Analysis version bcl2fastq2 v2.20 were run to demultiplex reads for all libraries and generate FASTQ files. The sequence data was run through the Slide- seqV2 pipeline with default settings^15, 16^. Code was retrieved from the Macosko Lab github page: https://github.com/MacoskoLab/slideseq-tools. Reads were mapped to the Sánchez Alvarado lab transcriptome and both mapped read 2 data and unmapped read 1 data were processed using Syrah, a Slide-seqV2 pipeline augmentation, in order to improve read count and reduce noise, resulting in a single digital gene expression matrix for each puck^44, 45^.

### Spatial embeddings

The R package Seurat v 4.0.1 was used to import the expression data for each puck and add slide x/y coordinates (the “slide” embedding)^18, 46–48^. Slide position and nUMI was used to determine which beads were under tissue fragments, label each fragment, and manually annotate the A-P axis for each fragment. The data were filtered to remove all beads except those under tissue, the data for all four pucks were combined, and the A-P angles were used to re-orient the fragments with the anterior at the top and the posterior at the bottom, organized into rows based on puck and timepoint (the “rows” embedding).

### Initial Analysis

Using the Seurat package in R^18^, the data were first normalized with SCTransform and the first 55 principal components were calculated. The PCA data were used to generate a UMAP embedding and to find clusters using FindNeighbors followed by FindClusters with resolution = 1. This resulted in 44 clusters.

### Label transfer for tissue annotation

Tissue type data from a recent scRNA-seq dataset were used to annotate the Slide- seqV2 data using Seurat’s FindTransferAnchors using 55 PCs, followed by TransferData^17, 18, 46^. The highest tissue label score for each bead was used as that bead’s tissue annotation.

### Spatial analysis of gene expression

Using the aligned “rows” embedding, the anterior (A) distance for each bead was calculated as the distance from that bead to the anterior-most bead in the same fragment. We determined which beads were on the edge of fragments by taking the centroid of each bead’s 60 closest neighbors. If the bead’s actual position was further than 30 units from the centroid, it was designated an edge bead. For all edge beads, edge distance was calculated as the distance to the closest edge bead within that same fragment (edge beads have edge distance zero).

### Gene cloning and riboprobe synthesis

Gene cloning was performed as previously described and plasmid constructs were transformed into E. coli strain HT115. For riboprobe synthesis, PCR products containing a single T7 promoter were amplified using the plasmid constructs as templates^17^. The primer sequences used are below. In vitro transcription was carried out at 25 μL scale according to the manufacturer’s instructions (Roche). Reaction products were precipitated by adding 80 μL ice cold ethanol, 12.5 μL ammonium acetate (7.5 M), and 2 μL glycogen (19-22 mg/mL, Sigma). Pellets were washed twice with 75% ethanol in water and resuspended in 100 μL deionized formamide (VWR).

PCR primers for riboprobe synthesis

PR244F: GGCCCCAAGGGGTTATGTG FM46: AGACCGGCAGATCTGATATCA

### *In situ* hybridization

Planarian fragments at relevant timepoints were fixed for single-color fluorescent *in situ* hybridization (FISH) using the NAC method and *in situ* hybridization was carried out as reported previously with the following customizations^43^. DAPI staining was performed at a final concentration of 2 ng/μL in MABT and incubated overnight at 4°C. Samples were cleared overnight in 20% ScaleA2 + DABCO solution and mounted the following day.

For double FISH, samples were fixed using NAFA fixation^49^. Samples were blocked in MABT containing 5% horse serum and 1% Western Blocking Reagent (Roche). *In situ* signals were developed as previously reported, but with the following customizations. Anti-Fluorescein-POD (Jackson Labs, Code 200-032-037) was applied in block solution at a concentration of 1:3000 overnight at room temperature. FAM tyramides were used to develop fluorescein probes. Peroxidase activity was inhibited using 100 mM sodium azide in PBS (+0.3% Tween-20) for 45 minutes on a shaker at room temperature. After washing > 6 times, Anti-Digoxigenin-POD (Roche, SKU 11207733910) was applied in block solution at a concentration of 1:1000 overnight at room temperature. A second signal was developed for the digoxigenin probes using rhodamine tyramides. DAPI staining was performed at a final concentration of 2 ng/μL in MABT and incubated overnight at 4°C. Samples were cleared overnight in 20% ScaleA2 + DABCO solution and mounted the following day.

For immunofluorescence of H3P, samples were fixed using NAFA fixation^49^. *In situ* signals were developed as above. Following development, samples were blocked, stained, washed, cleared as described. Phophorylated Histone H3 was visualized with the following antibodies: Abcam Rabbit Anti-Phospho-Histone H3 (S10+T11) (ab32107), Abcam Goat anti-Rabbit IgG H&L (Alexa 647) (ab150079).

### Microscopy

Automated imaging was used to capture 10X magnification confocal microscopy images of fluorescent *in situ* hybridizations. Overview images of slides were acquired with a Plan Apochromat Lambda 4X objective lens (N.A. 0.2, 1.735 μm/pixel) and masked using a Fiji macro in a Nikon Elements job. DAPI fluorescence was used to identify planarian fragments. Final images were acquired with a Plan Apochromat Lambda 10X objective lens (N.A. 0.45, 0.78 μm/pixel). Images were batch stitched using Fiji macros (https://github.com/jouyun/smc-macros).

High magnification confocal images of *mmp-1*^+^ cells were acquired with an Orca Flash 4.0 sCMOS 100 fps camera at full resolution on a Nikon Eclipse Ti2 microscope equipped with a Yokogawa CSU W1 10,000 rpm Spinning Disk Confocal with 50 μm pinholes. Samples were illuminated with 405 nm (3.9 mW), 488 nm (8.5 mW), and 561 nm (6.1 mW) lasers (LUNV 6-line Laser Launch) with nominal power measures at the objective focal plane. This spinning disk confocal is equipped with a quad filter for excitation with 405/488/561/640 nm. Emission filters used to acquire this image were 430-480 nm, 507-543 nm, and 579-631 nm. The *mmp-1^+^* cells near the wound site were identified and used to define upper and lower bounds for confocal imaging. A stack of images was acquired with 0.5 μm step size between each image. A Nikon Plan Apochromat 100X oil objective lens (N.A. 1.49, 0.065 μm/pixel) was used to acquire the image. Selected images were cropped in Fiji and used for 3D rendering.

### Image Analysis

“A-distance” values were quantitated as follows: tail sections were selected from multiple rounds of automated 10X imaging on the spinning disc, as described above. A single z-slice was chosen that corresponds to the main plane of gene localization. First, the tail slice image was rotated to place the wound site in a consistent position. Tail slices were of similar but not identical size. To allow for averaging, images were scaled to the same size. To remove background, a rolling ball background subtraction of 200 pixels was applied, followed by a Gaussian blur of radius 50 pixels. A line profile was drawn starting at the wound site to the end of the tail slice, and a kymograph was obtained averaging over the width of the tail slice. This provided a distribution of intensity of the FISH signal, anterior to posterior. The kymographs for a given FISH probe were averaged to obtain the average A-distance plots. Distributions of fluorescence intensity were compared using the Kolmogorov-Smirnov test.

### Spatial analysis of imaging data

1 mm spinning disk confocal stacks were projected to a smaller series of 6 mm stacks, where the non-DAPI channels were maximally projected in order to collect all potential signal from one cell into a single image and the DAPI projected only the middle two sections. Projecting the entire DAPI over this range had negative effects on the results. Segmentation was then run on the projected DAPI channels using the ’cyto’ model from Cellpose 2.0 with a diameter of 50 pixels^24, 25^. Using the nuclear segmentation masks, the integrated signal intensities for all channels were calculated. Signal intensities for each nucleus are normalized by finding the 30th percentile of intensities for each channel/slice/file and dividing all intensities at the same channel/slice/file by that value. This accounts for signal attenuation at different penetration depths and for animal-to-animal variability. A threshold is then selected based on visual inspection and then applied to determine which nuclei are positive for the various labels. Hecatonoblasts were segmented using thresholds, and a distance map tabulated to generate the distance data. For the intestine, segmentation was performed manually in Fiji and a distance map similarly generated. A github repository contains the jupyter notebooks used to perform the analysis: https://github.com/jouyun/Mann_2023.

### 3D rendering of *in situ* confocal microscopy

3D renderings and movies were made using Fiji 3D Viewer^50, 51^ with a downsample of 4 and a threshold of 2, 20, and 20 for *mmp-1, piwi-1*, and DAPI signal, respectively.

Resulting surfaces were exported as wavefront files. Meshes were imported into Blender (BO Community) for still images and movies.

## DATA AVAILABILITY

The Slide-seqV2 data generated by this study have been deposited in GEO at GSE199348. Data will be made available upon publication. Bead metadata are available in Supplementary Table 5.

## CODE AVAILABILITY

Relevant scripts generated by this study have been deposited to GitHub. Scripts for analysis of Slide-seqV2 data are available at: https://github.com/0x644BE25/smedSlide-seq Jupyter notebooks for spatial analysis of imaging data are available at: https://github.com/jouyun/Mann_2023

## ACKNOWLEDGEMENTS

We thank Fei Chen and Evan Murray for providing spatially barcoded pucks and experimental support, the Stowers Planaria Core for support with animal husbandry, Yongfu Wang for assistance with tissue sectioning, Michael Peterson for assistance with Illumina sequencing, Madelaine Gogol and Hua Li for bioinformatic support, Jay Unruh, and members of the Sánchez Alvarado lab for helpful discussions and feedback. A.S.A. is an investigator of the Howard Hughes Medical Institute (HHMI) and the Stowers Institute for Medical Research. B.W.B.-P was a Jane Coffin Childs Memorial Fund Postdoctoral Fellow. F.G.M. is an HHMI Postdoctoral Fellow. This work was supported in part by NIH R37GM057260 to A.S.A.

## AUTHOR CONTRIBUTIONS

B.W.B.-P. and F.G.M. designed the experiment. B.W.B.-P., F.G.M., S.M., and J.A.M. created tissue sections for analysis. K.E.H. and A.P. performed library preparation and sequencing. B.W.B.-P., F.G.M., C.E.B., E.R.D., D.M.V., S.C., S.A.M, B.D.S. and A.S.A. analyzed the data. B.W.B.-P., F.G.M., E.R.D., D.M.V., and L.E.M. performed and imaged in situ hybridizations. C.G.H. optimized in situ hybridizations on regenerating planarians. S.H.N. rendered light microscopy data into three-dimensional models.

---

**Fig. S1:**
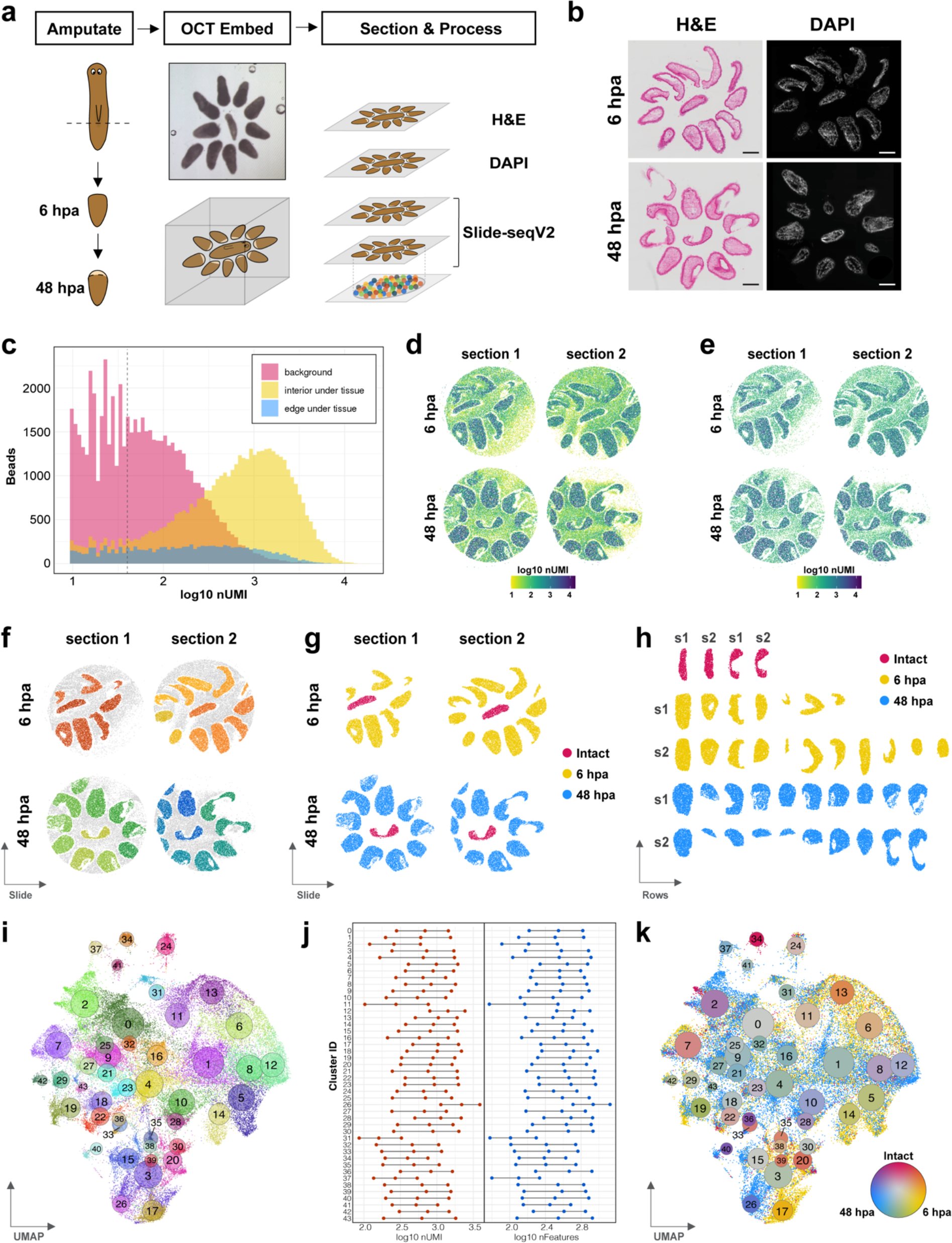
A spatial atlas of planarian regeneration. **a,** Diagram of experiment. Approximately 10 regenerating planarian tail fragments at both 6 and 48 hpa were co-embedded around a single intact planarian and sectioned at a thickness of 10 microns for Slide-seqV2. Two adjacent sections were analyzed on separate pucks. Two additional sections were stained and imaged. **b,** Images of adjacent sections from both timepoints stained with H&E or DAPI. Scale bars = 200 microns. **c,** UMI distributions of beads under planarian tissue (yellow), not under tissue (pink), or around the edges of planarian fragments (blue). **d,** Spatial visualization of UMI counts on pucks using default 10 UMI cutoff for Slide-seqV2. **e,** Spatial visualization of UMI counts on pucks using 40 UMI cutoff for Slide-seqV2. **f,** Manual annotation of beads under each planarian fragment. **g,** Beads under fragments colored by timepoint of origin. **h,** “Row-embedding” of beads under the regenerating tissue fragments. Colors indicate timepoint of origin. Fragments have been rotated with anterior poles facing upward. **i,** UMAP plot of all under-tissue beads in the dataset, colored by cluster number. Area of the circles is proportional to the number of beads in the cluster. **j,** Plot of UMI counts (left, red) and number of features (right, blue) for all beads in each cluster. Dots represent the first quartile, median, and third quartile for each cluster. **k,** UMAP plot of complete dataset. Clusters are labeled with circles proportional in area to the number of beads and colored by relative proportion of beads from each timepoint.

**Fig. S2:**
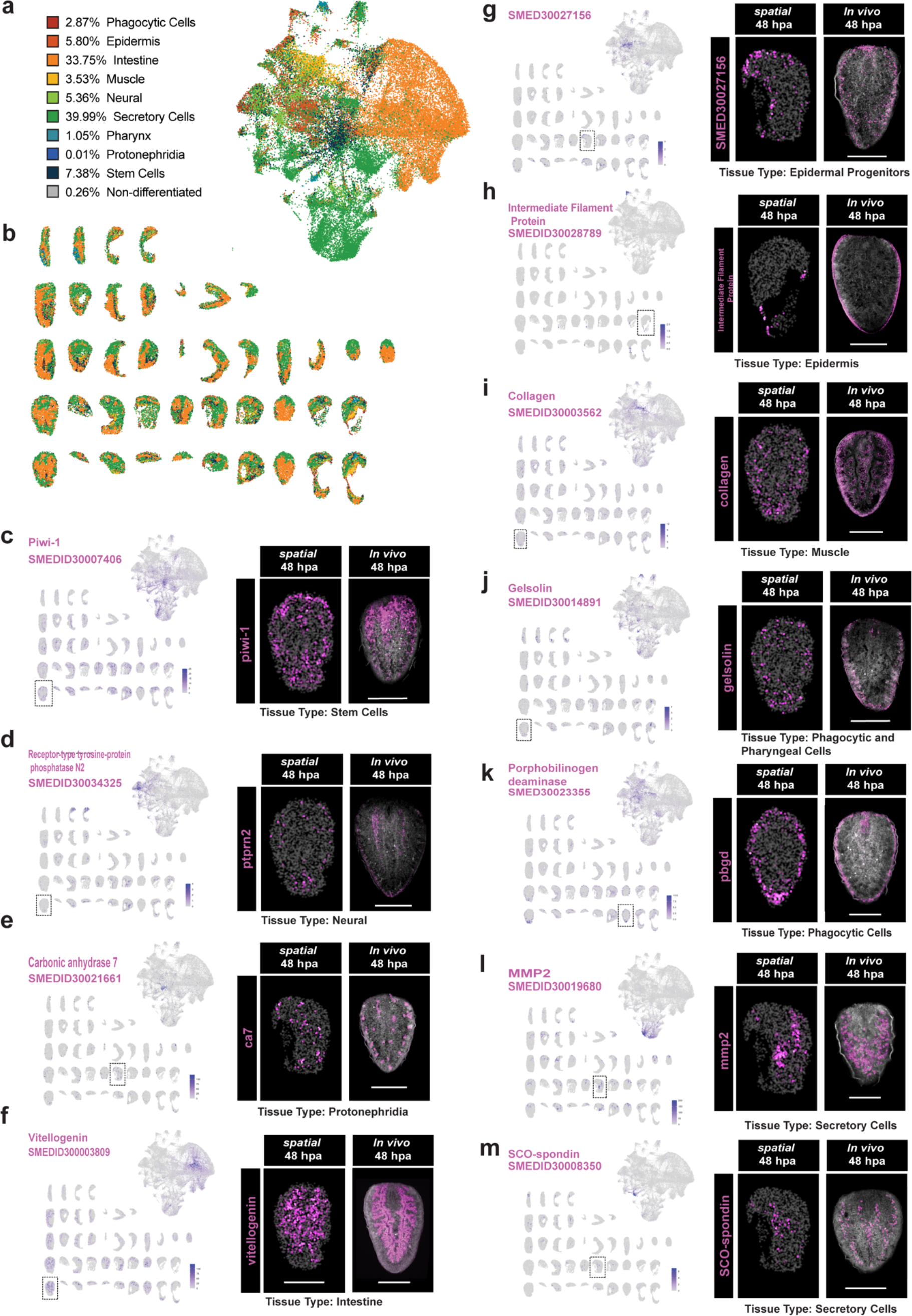
Slide-seqV2 data recapitulates expression patterns *in vivo*. **a,** UMAP plot of all under-tissue beads colored by dominant tissue signature. **b,** ‘Row embedding’ of beads, colored by dominant tissue signature. Anterior = up for all fragments. **c-m,** Comparison of Slide-seqV2 and *in situ* data for 11 markers of various tissues. For each, left: UMAP plot and row embedding showing gene^+^ beads in Slide- seqV2. Color intensity is proportional to expression level. Each gene is scaled separately. Right: Slide-seqV2 data for boxed fragment compared to *in situ* hybridization against that gene. Scale bars = 500 microns.

**Fig. S3:**
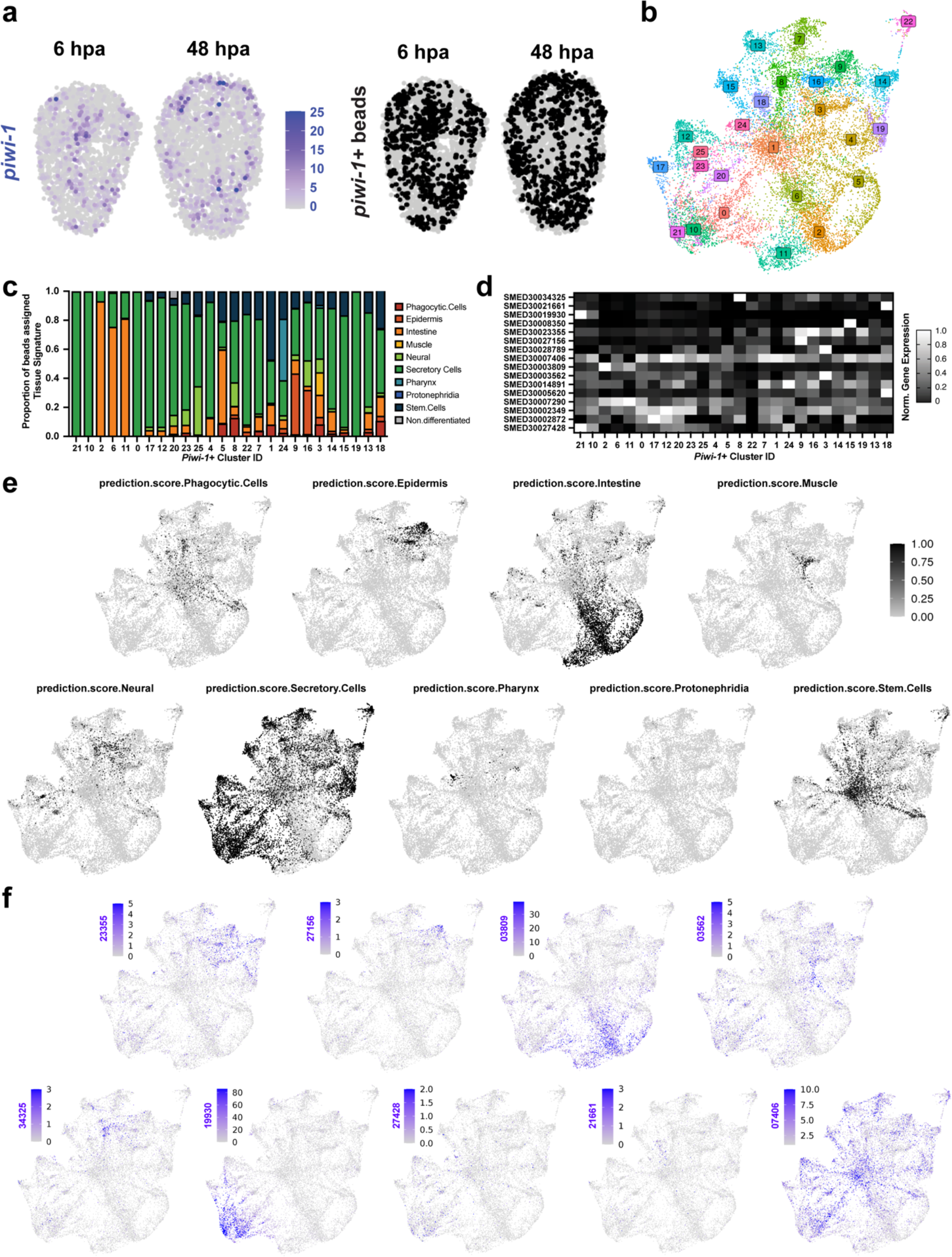
p*i*wi*-1^+^* bead subset is dominated by secretory and intestinal signatures. **a,** *piwi-1* expression captured by Slide-seqV2. Left: number of UMI for *piwi-1* reads by timepoint for example fragments. Right beads with at least one *piwi-1* read. **b,** UMAP plot of *piwi-1^+^* bead subset colored by cluster. Cluster numbers are superimposed on the plot. **c,** Plot showing top tissue annotation assigned by Seurat’s LabelTransfer function for each bead in a given *piwi-1^+^* cluster. **d,** Heatmap showing expression of tissue markers for each *piwi-1* cluster. Rows are individually scaled relative to each marker’s maximum expression level. **e,** UMAP plots of *piwi-1*^+^ bead subset shaded by LabelTransfer tissue annotation score for each of the nine major tissues profiled in Benham-Pyle *et al.*, 2021. **f,** UMAP plots of *piwi-1*^+^ bead subset shaded by UMI count for markers of each of the nine major tissues from **e.** Leading “SMED300” was dropped from gene names to save space.

**Fig. S4:**
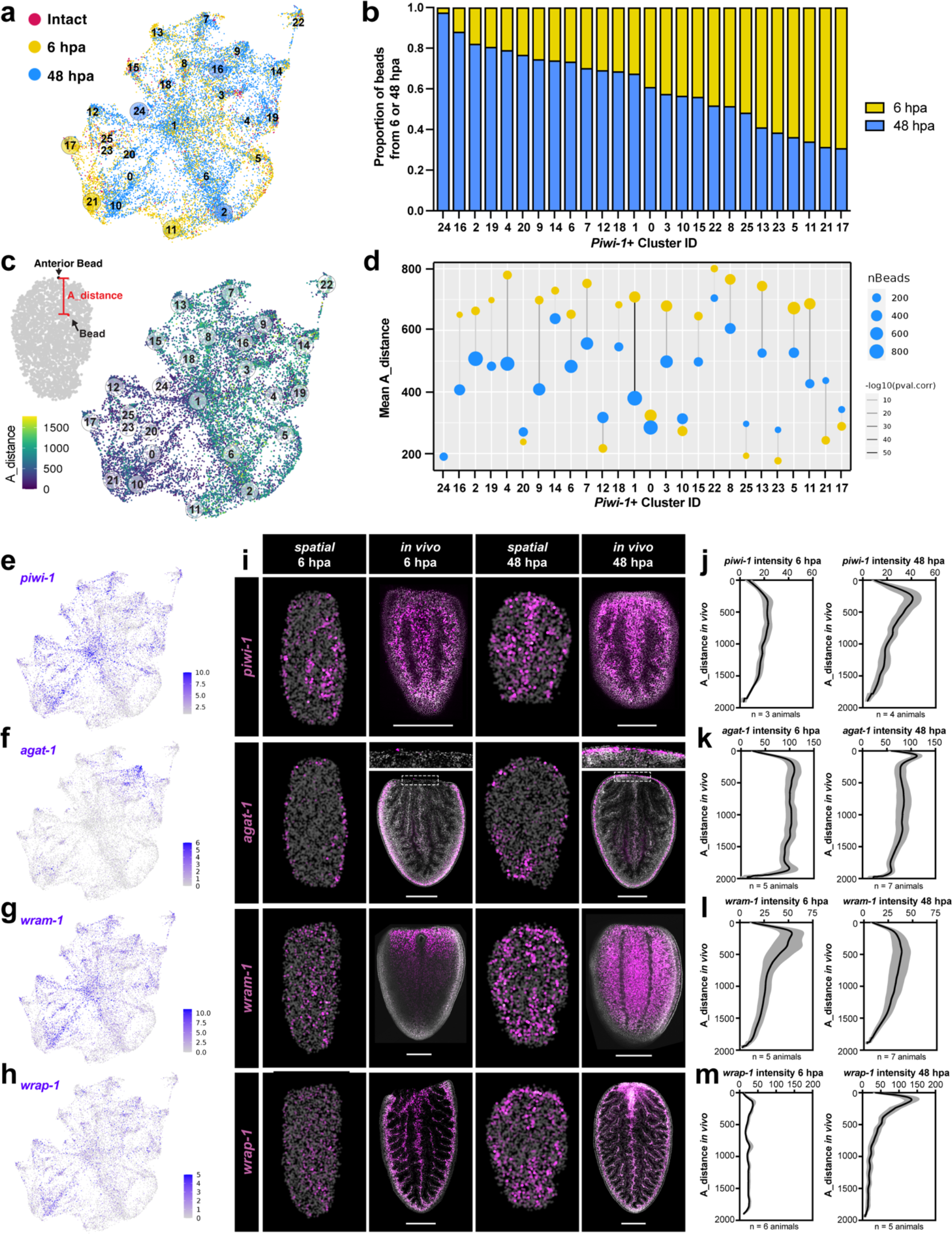
Identification of local and global transcriptional responses to wounding. **a,** UMAP plot of *piwi-1*^+^ bead subset colored by timepoint of origin. Cluster numbers are superimposed on the plot. **b,** Plot showing timepoint of origin for all beads in a given *piwi-1*^+^ cluster. **c,** Left, schematic of ‘A-distance’ calculation. Right, UMAP plot of *piwi-1^+^* bead subset colored by A-distance. Cluster numbers are superimposed on the plot. **d,** Plot showing change in A-distance over time. For each cluster, yellow dot indicates average A-distance at 6 hpa, blue dot indicates average A-distance at 48 hpa. Dot size is proportional to number of beads. **e-h,** UMAP plots of *piwi-1^+^* bead subset shaded by UMI count for genes with notable A-distance patterns. **i**, Comparison of expression patterns captured by Slide-seqV2 and by *in situ* hybridization. Third row: *wound- responsive associated with muscle 1* (*wram-1*). Fourth row: *wound responsive associated with piwi-1* (*wrap-*1). Scale bars = 500 um. **j-m,** Plot depicts distributions of *in situ* hybridization signals for A-distance genes. Y-axis is distance from wound site.

**Fig. S5:**
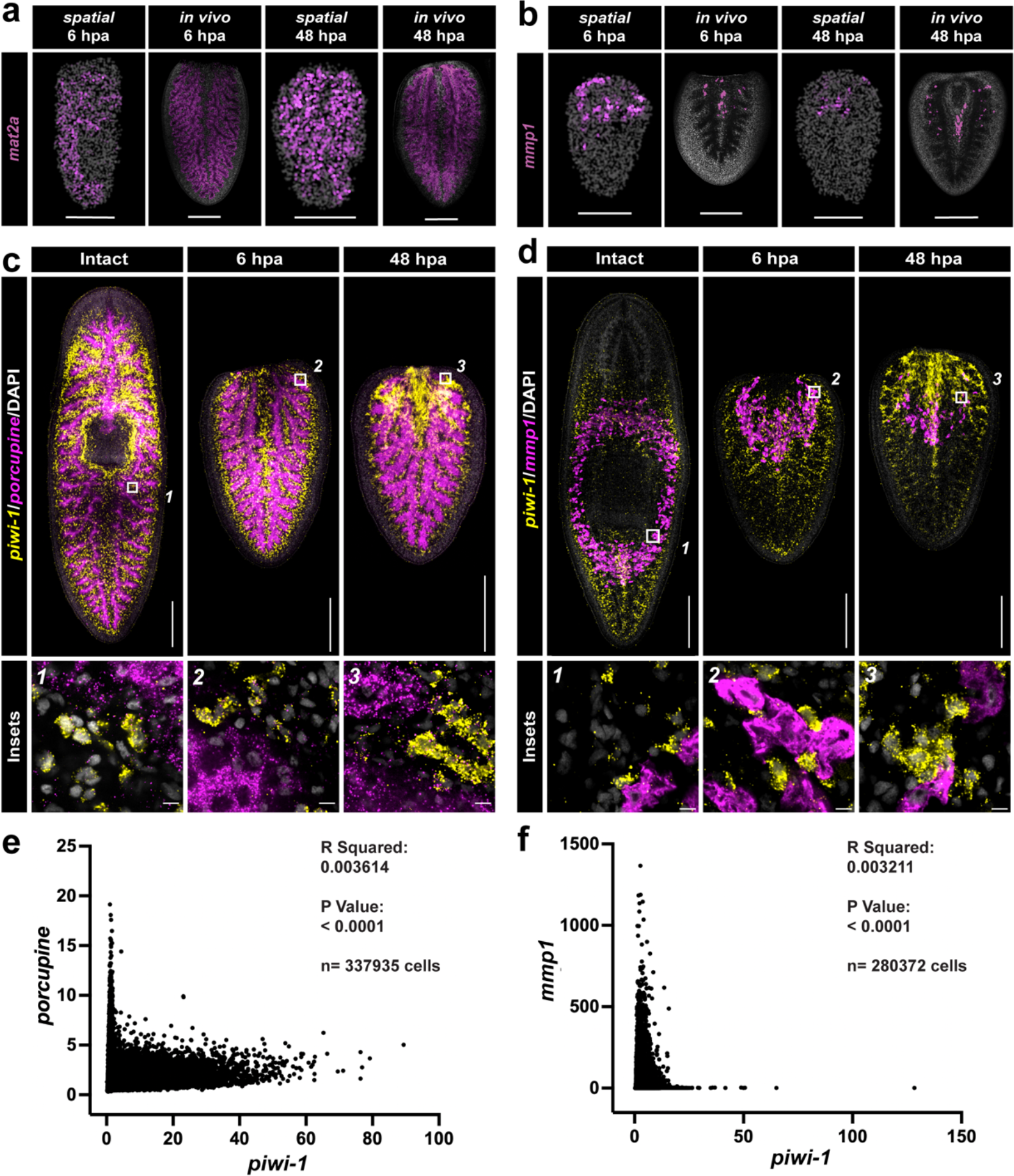
Beads reflect close associations between cell types. **a,** Comparison of *mat2a* expression patterns via Slide-seqV2 (first and third panels) and fluorescent *in situ* hybridization (second and fourth panels) maximum intensity projections. Scale bars = 500 um. **b,** Comparison of *mmp-1* expression patterns via Slide-seqV2 (first and third panels) and fluorescent *in situ* hybridization (second and fourth panels) maximum intensity projections. Scale bars = 500 um. **c,** Double fluorescent *in situ* hybridization of the intestinal marker *porcupine* (magenta), *piwi-1* (yellow), and DAPI (white) in a single confocal Z-section. Areas of interest are indicated by white boxes and shown in greater detail as insets below. Upper scale bars = 500 um, lower scale bars = 10 um. **d,** Double fluorescent *in situ* hybridization of the intestinal marker *mmp-1* (magenta), *piwi-1* (yellow), and DAPI (white) in a single confocal Z- section. Areas of interest are indicated by white boxes and shown in greater detail as insets below. Upper scale bars = 500 um, lower scale bars = 10 um. **e,** Plot of fluorescence intensity of *piwi-1* versus *porcupine* in 337,935 cells produced by CellPose analysis of confocal image stacks. **f,** Plot of fluorescence intensity of *piwi-1* versus *mmp-1* in 280,372 cells produced by CellPose analysis of confocal image stacks.

**Fig. S6:**
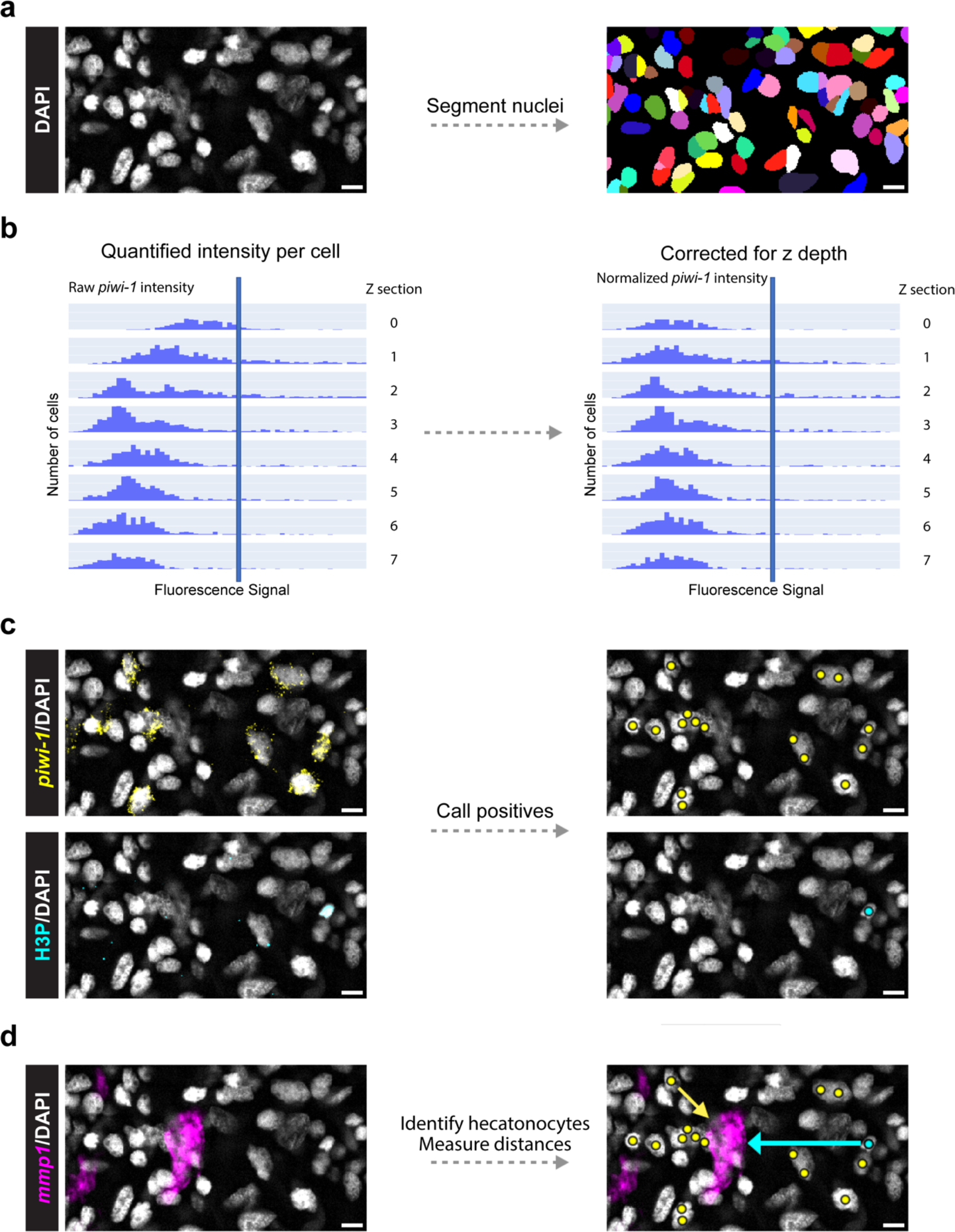
Overview of CellPose analysis workflow. **a,** CellPose identifies cells in confocal image stacks by using DAPI signal to segment nuclei^24, 25^. Scale bars = 10 microns. **b,** Fluorescence intensity is quantified for each channel in a segmented area. Data are normalized to correct for signal attenuation as a function of depth. **c,** Positive cells are called using a threshold fluorescence intensity. **d,** Calculating minimum distance between cells of interest and hecatonoblasts or the intestine.

**Fig. S7:**
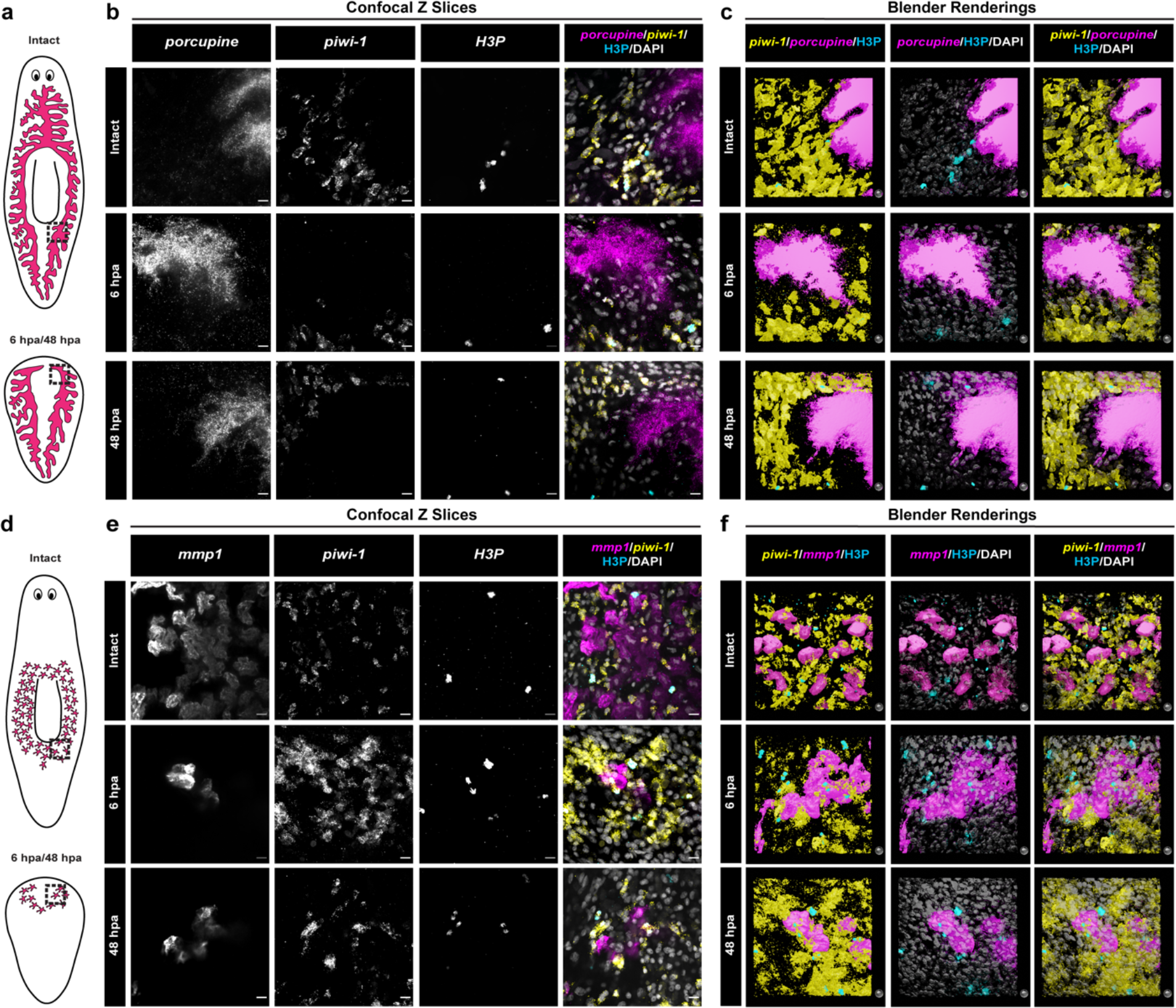
Individual fluorescent channels and 3D renderings for colocalization analysis. **a,** Cartoon of the intestine (magenta) in intact (top) and regenerating (bottom) planarians. **b,** Individual fluorescent channels and 3D renderings for single-slice merged confocal images shown in Figure 2. First through Fourth columns: individual fluorescent channels for *porcupine*, *piwi-1*, H3P, and a merge, respectively. Fifth through Seventh columns: 3D renderings of confocal image stacks shown for viewability without DAPI, without *piwi-1*, and with all four channels, respectively. Scale bars and spheres = 10 microns. **c,** Cartoon of hecatonoblasts (magenta) in intact (top) and regenerating (bottom) planarians. **d,** Individual fluorescent channels and 3D renderings for single- slice merged confocal images shown in Figure 3. First through Fourth columns: individual fluorescent channels for *mmp-1*, *piwi-1*, H3P, and a merge, respectively. Fifth through Seventh columns: 3D renderings of confocal image stacks shown for viewability without DAPI, without *piwi-1*, and with all four channels, respectively. Scale bars and spheres = 10 microns.

**Fig. S8:**
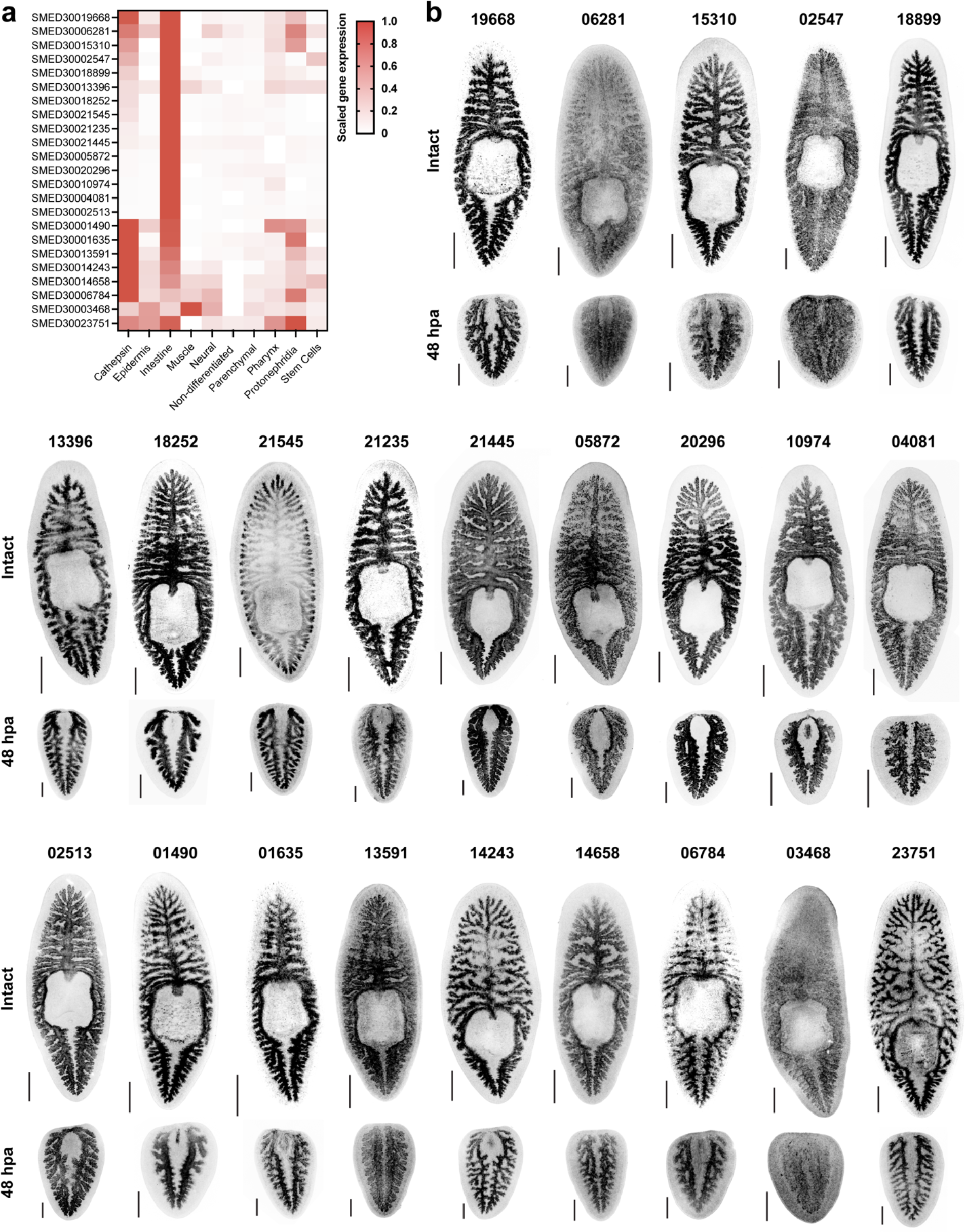
Expression patterns of intestinal gene set. **a,** Heatmap shows expression levels of intestinal genes by tissue in a single-cell RNA- seq regeneration timecourse^17^. **b,** Expression patterns of intestinal genes in intact (upper) and 48 hpa regenerating fragments (lower) visualized by *in situ* hybridization. Leading “SMED300” was dropped from gene names to save space. Scale bars = 500 microns.

**Fig. S9:**
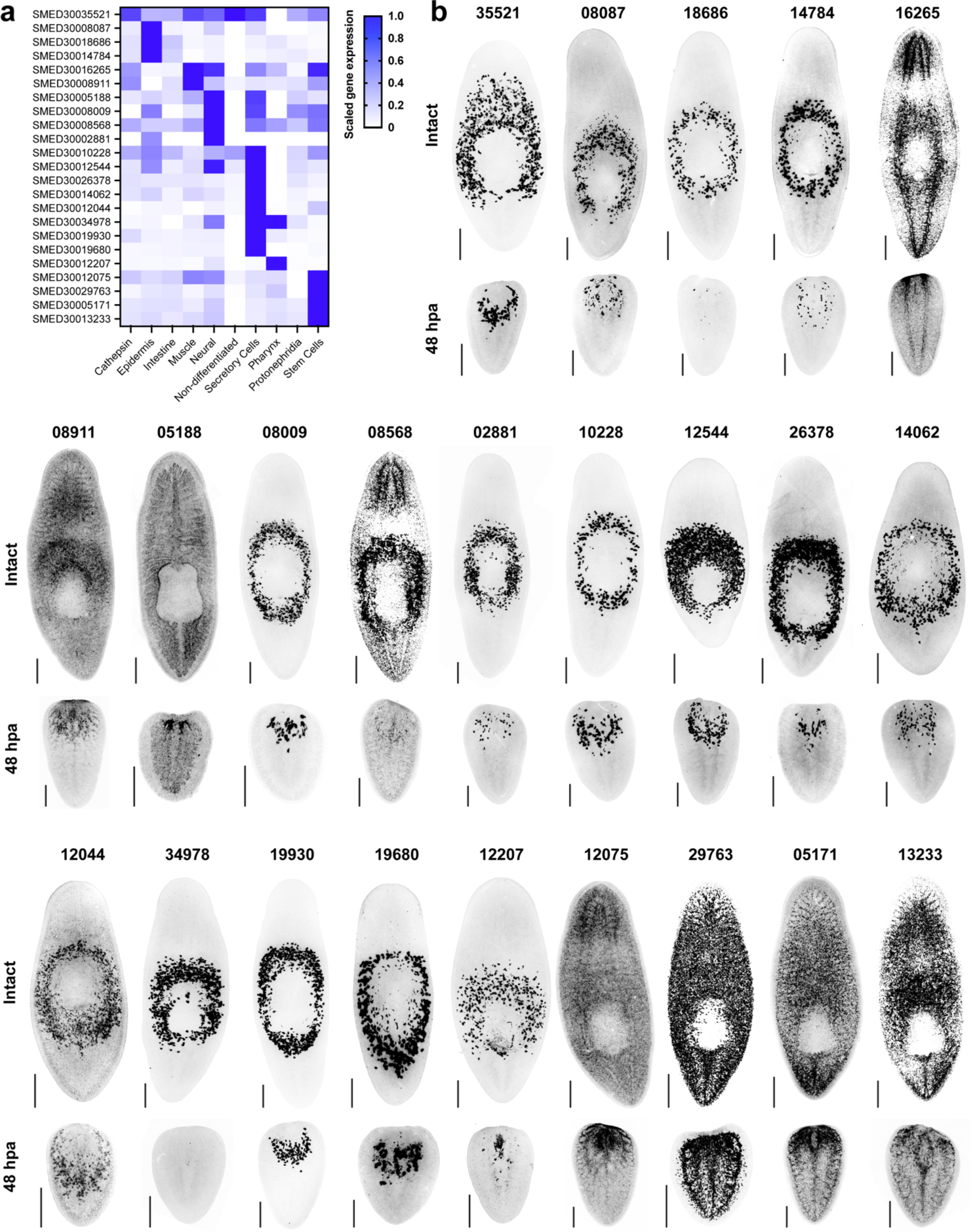
Expression patterns of secretory cell gene set. **a,** Heatmap shows expression levels of secretory cell genes by tissue in a single-cell RNA-seq regeneration timecourse^17^. **b,** Expression patterns of intestinal genes in intact (upper) and 48 hpa regenerating fragments (lower) visualized by *in situ* hybridization. Leading “SMED300” was dropped from gene names to save space. Scale bars = 500 microns.

**Fig. S10:**
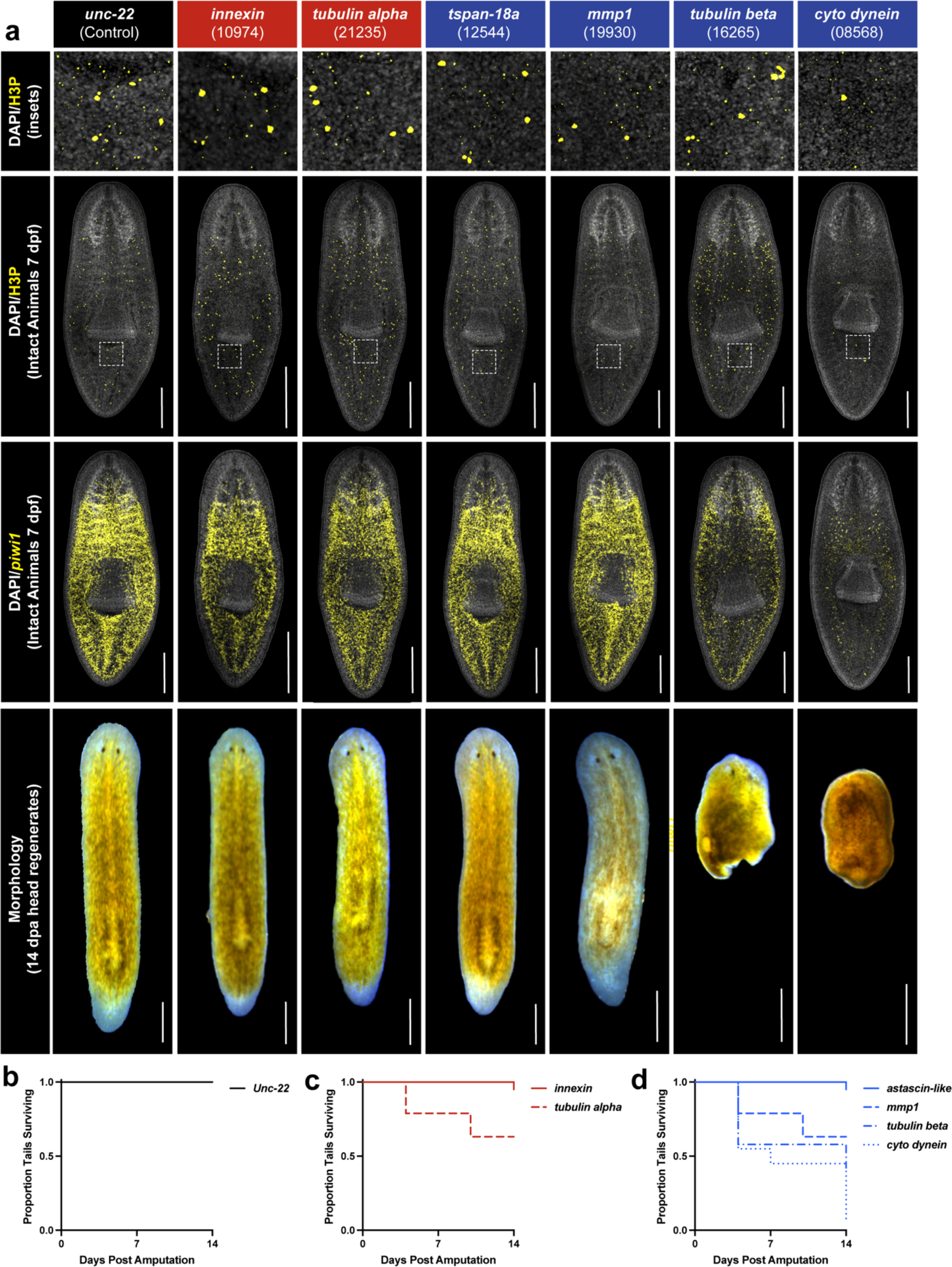
The intestine and hecatonoblasts promote proper regeneration by regulating stem cell proliferation and accumulation. **a**, Top row: Detail of confocal max projections of mitotic H3P+ cells (yellow) with DAPI (white) in intact RNAi animals. Second row: Confocal max projections providing overview of intact, uncut animals. Boxed areas indicate regions of interest in top row. Third row: Confocal max projection of intact animals stained by *in situ* hybridization against *piwi-1*+ (yellow) and DAPI (white) seven days post feeding RNAi food (7 dpf). Fourth row, Regenerated planarian fragments with new tails 14 dpa. Gene names boxed in red are intestinal; those boxed in blue are secretory. Scale bars = 500 um. b, Survival curve of *unc-22* control animals. c, Survival curves of two intestinal RNAi conditions. d, Survival curves of five secretory RNAi conditions.

## TABLES

Supplementary Table 1: Marker genes identified from Slide-seqV2

Supplementary Table 2: Statistical information for spatial analysis of images

Supplementary Table 3: Statistical information for mitotic density

Supplementary Table 4: Planarian gene nomenclature

Supplementary Table 5: Slide-seqV2 metadata

